# Novel Function of Transcription Factor TTF1 in UV Mediated DNA Damage Repair in Mammalian Cells

**DOI:** 10.1101/2023.05.09.540020

**Authors:** Kumud Tiwari, Snehendu Bose, Neelu Mishra, Samarendra Kumar Singh

**Affiliations:** School of Biotechnology, Institute of Science, Banaras Hindu University, Varanasi, Uttar Pradesh, 221005, India

## Abstract

Various DNA repair machineries have evolved in the cell to maintain the integrity of the genome for proper functioning of the same. Repair of ultraviolet (UV) irradiation mediated DNA damage occurs by nucleotide excision repair pathway through transcription coupled repair (TCR), a process in which the damage is repaired on transcriptionally active genic regions. This process requires various protein complexes including heterodimer DNA Damage Binding 1 (DDB1) protein. Defects in TCR have been found in patients with mutations in the Cockayne syndrome (CS) group A and group B genes and in the Xeroderma pigmentosum (XP) group G gene. Transcription factors (TFs) play a very important role in regulation of TCR system, especially those TFs which binds to DNA at specific loci. Several TFs have been shown to modulate the repair of photolesions either by inducing or inhibiting the TCR. However, the mechanism behind their action is not very clear. Mammalian TTF1 is an essential multifunctional transcription factor involved in transcription initiation, termination, DNA fork blockage, chromatin remodelling etc., and has been shown to interact with CSB protein. Hence, to discover its role in TCR and to identify its interaction partners, we purified this protein and did a pull-down assay with HEK293T cell lysate followed by LC-MS and discovered DDB1 as one of its major interactors. This established our confidence that TTF1 protein might be playing a critical role in TCR. Further, we discovered that, upon UV mediated DNA damage in HEK293T cells the expression of TTF1 is significantly induced and is co-localized with _γ_H2AX protein. To our surprise, we found that after knockdown of TTF1, DDB1 level decreases in HEK293T cells while knockdown of DDB1, increases TTF1 level in the cells. Hence, our study opens up a new avenue towards exploring a noble function of the transcription factor TTF1, which in turn could establish the potential to develop therapeutics towards cancers and other diseases.

## Introduction

In order to maintain over all health, various DNA damage repair mechanisms exist within the cells. Damage caused by irradiation especially UV rays is repaired by nucleotide excision repair (NER) mechanism. The two pathways of NER are transcription coupled NER (TC-NER) and global genomic NER (GG-NER). Mis-regulation or deficiency in NER has been reported to cause many fatal abnormalities, among them is disorders like Cockayne syndrome and Xeroderma pigmentosum [1–4]. GG-NER repairs and restores damaged DNA strands all across the genome. As a part of the damaged DNA binding protein complex (UV-DDB), which senses and binds to UV-damaged DNA, DNA binding protein 1 (DDB1) is a conserved protein that participates in the global genomic NER. DDB1 (also known as p127) and DDB2 (also known as p48) are two subunits of the complex UV-DDB. Despite being majorly involved in UV mediated NER, cell proliferation and apoptosis, it is also involved in many cellular processes as it interacts with various other multifunctional proteins. DDB1 which primarily is a cytoplasmic protein, diffuses into the nucleus following UV exposure and complexes with DDB2 or Cockayne Syndrome Protein (CSA) [5–7] in response to DNA damage. In contrast to the DDB1/CSA complex, which scans the anti-sense strand of transcribed genes (TC-NER), the DDB1/DDB2 complex facilitates the detection of photolesions throughout global genome repair (GG-NER). DDB1 is a critical part of Cul4A-RING ubiquitin E3-ligases (CRL4) serving as an adapter protein for Cullin 4A (Cul4A) and CUL4-associated components (DCAFs), interacts with many WD40 proteins as a receptor substrate and aids in the formation of more than 90 E3-ligase complexes (Figure 1)[8]. DDB1-Cul4A ubiquitin-proteasome system ubiquitinates and degrades many important factors like p27, p21, cdt1 etc. The discovery that UV-DDB is a part of an E3-ubiquitin ligase complex raises the possibility that ubiquitination and DNA damage sensing and repair through NER are connected [9]. In fact, the CRL^DDB1^ complex has a role in both the onset of GG-NER and activation of transcription during TC-NER [5,6,10].

**Figure 1:**
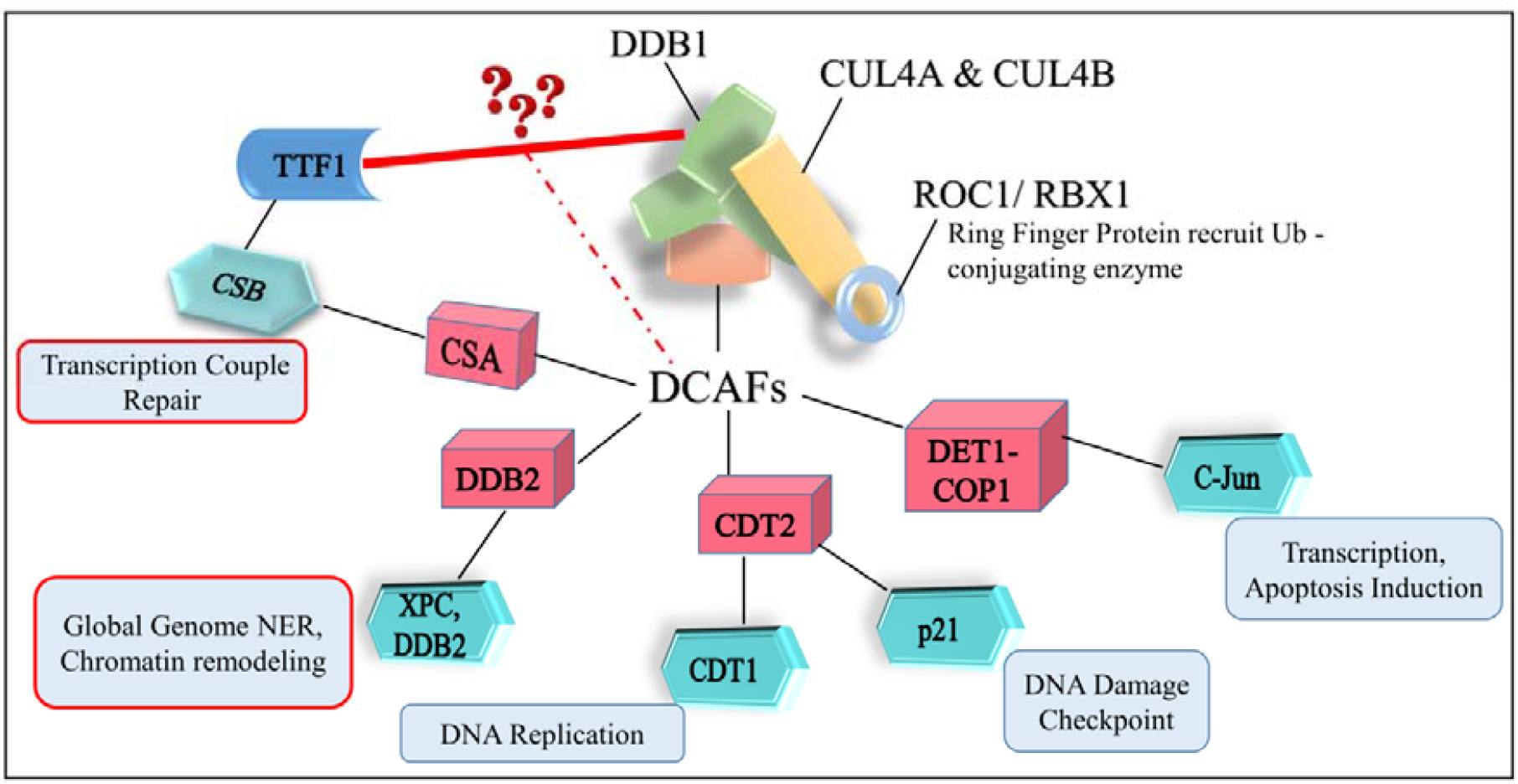
Molecular architecture of CUL4 Ligase with DCAFs interacting proteins and their respective function/s with hypothesis that what would the role of TTF1 in this cascade.

Transcription Termination Factor (TTF1) is a multifunctional protein engaged in various essential cellular functions like transcription initiation and termination, replication polar fork blockage, chromatin remodelling, cell cycle regulation etc. Towards executing the above processes, it interacts with numerous regulatory proteins, including TTF1 Interacting Protein (TIP 5), Pol I and transcript release factor (PTRF), Mouse Double Minute 2 (MDM2), Cockayne Syndrome B (CSB), Alternative Reading Frame (ARF), etc. (Table 1) and thereby catalyzes critical functions in mammalian cells. These diverse functions make TTF1 an essential protein for the survival of the cell. Hence, mis-regulation of TTF1 could lead to various cellular defect leading to oncogenic transformation of the cells. Overexpression of TTF1 has been reported in various tumours, indicating that TTF1 is required in higher quantities to meet the higher rate of ribosome biogenesis to cope up with hyper-proliferation of tumorous cells. Considering the essential and multifunctional nature of this protein, it becomes important than ever to characterize the mechanism behind its function and regulation [11–13].

**Table 1:**
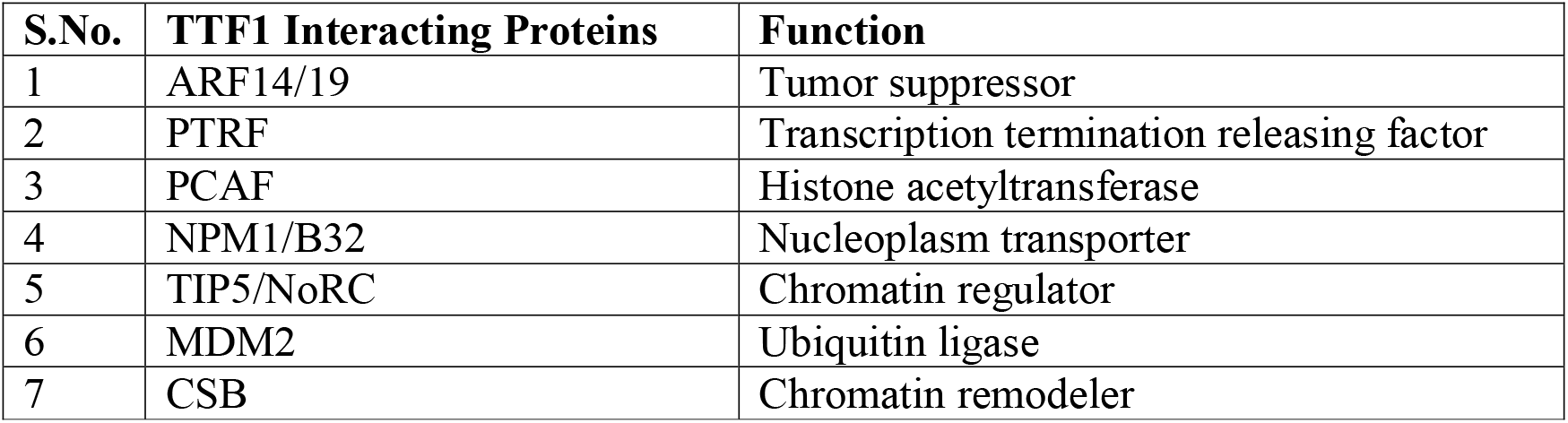
TTF1 interacting proteins.

In this study, we report the discovery of TTF1’s noble interactor DDB1. We found out that suppression of TTF1 downregulates DDB1 expression while on the contrary DDB1 suppression upregulates TTF1. Upon UV induced DNA damage in the HEK293T mammalian cells, TTF1 is being accumulated at the damage sites co-localized with DNA damage marker _γ_H2AX[14]. This is the first ever study to show the role of TTF1 in UV mediated DNA damage response (and probably in repair as well).

## Materials and Methods

### 1. Cell culture

HEK293T (Human kidney embryonic 293T) cells (NCCS, Pune, India) were cultured in Dulbecco’s modified Eagle’s medium (DMEM) containing 10% Fetal bovine serum (FBS) (Gibco, USA). 1% Pen-Strep (Penicillin-Streptomycin, Gibco, USA) was added to culture media as routine practice. The cells were grown in incubators with 5% CO_2_ and 95% humidity at 37°C.

### 2. Pull down assay followed by Mass spectrometry

Histidine tagged codon optimized mammalian TTF1 was expressed in BL21-DE3 cells and purified as mentioned in Tiwari et. al [12]. This purified protein was then immobilized on Ni-NTA column. Subsequently, HEK293T cells were lysed (in lysis buffer; 25 mM Tris pH 7.5, 500 mM KCl, 10% glycerol, 9 mM BME + protease inhibitor cocktail), followed by sonication and centrifugation at 15K rpm, 4°C for 15 min for obtaining clear lysate. The lysates from HEK293T cells were subjected to Ni-NTA beads in the presence or absence of immobilized TTF1 (in above mentioned binding buffer). Eventually, the Ni-NTA beads were then washed with binding buffer (20 column volume) and eluates were sent for mass spectrometry analysis.

### 3. siRNA transfection

Small interfering RNAs (siRNAs) were used at a final concentration of 50 nM. 0.15 × 10^6^ logarithmically growing HEK293T cells were seeded in each well in a 6 well plate. Subsequently, cells were transfected with siDDB1, siTTF1 and scramble (siRNA non-target) using turbofect™ (Thermo Fisher Scientific, Massachusetts, USA) transfection Reagent. Cells were then cultured as mentioned above. After 24 hrs of transfection, the medium was replaced with fresh medium containing 10% FBS, and 48 hrs later the cells were harvested for further procedures. siDDB1 and siTTF1 were purchased from Helix technology (India).

### 4. UV irradiation

A total of 0.2 × 10^6^ HEK293T cells were seeded onto 30 mm plates in 3 ml of fresh DMEM medium. After incubation for 24 hrs as mentioned above, cells were washed and covered with 1 ml of phosphate-buffered saline (PBS), and the monolayer of sub-confluent cells were irradiated with UV-C (200-290 nm) having energy flux of 30 J/m^2^. The PBS was then replaced with 3 ml of fresh DMEM medium and re-incubated the cells for the indicated period of time.

### 5. Western Blot Analysis

HEK293T cell were harvested and washed with PBS after 5 mins, 10 mins and 15 mins post UV-C irradiation as mentioned above. Further procedure was performed as described by Garima et. al [15]. TTF1 and DDB1 primary antibodies were purchased from Abcam (Cambridge, UK) and Cell Signalling Technology (Massachusetts, USA) respectively.

### 6. Co-immunoprecipitation

HEK293T cells were lysed at 4°C for 1 hr in a lysis buffer (50 mM Tris-Cl [pH-7.5], 10% glycerol, 100 mM NaCl, 1 mM EDTA, 1 mM DTT, 0.1% TritonX-100, 1 mM PMSF, Protease inhibitor cocktail) followed by sonication and then centrifugation (15,000 rpm, 4°C for 15 min). The cell lysate was then mixed with the respective IgG immobilized with agarose A beads and incubated for 1 hour at 4°C with rotation. Beads were then washed 3 times with 1 mL lysis buffer to remove non-specific interactors. The proteins were then eluted in 2X volume lammeli buffer followed by heating at 98°C to release the bound protein complex. The sample was then centrifuged at 8K rpm for 5 mins and the supernatant was resolved on SDS-PAGE (8 %) followed by western blotting with respective antibodies as mentioned above.

### 7. Immunofluorescence and Confocal microscopy

0.2 × 10^6^ HEK293T cells were seeded on sterilized coverslips. After 24 h, the cells were locally irradiated and re-incubated as mentioned above. The cells on coverslips were then washed gently in PBS, and fixed in buffer containing freshly made 4% paraformaldehyde for 20 min. The cells were then permeabilized using 0.1% Triton X-100 in PBS (PBST) for 15 min at 4°C. Thereafter, the samples were washed with PBS and blocked for 1 hr in PBS containing 5% bovine serum albumin (BSA). Cells were then incubated overnight in dark with specific primary antibodies: rabbit polyclonal anti-TTF1 (1:100), rabbit polyclonal anti-_γ_H2A.X (1:100, CST, Massachusetts, USA) followed by secondary antibody at 4°C (in dark humid chamber). They were then washed with PBS and mounted on slide using mounting media (Vectashield, Vector Laboratories, San Francisco, USA). Images were acquired on confocal microscope (Leica, Wetzlar, Germany) using a 20X (magnification) and 1.4 (aperture) objective with identical settings to ensure comparability.

## Results

### 1. DDB1 protein is a potential interactor of TTF1

After subjecting the HEK293T cell lysate onto the immobilized TTF1 protein, we performed mass spectrometry of the pull-down fraction to identify the TTF1 interacting proteins (Figure 2A). DDB1 which is an essential protein involved in DNA damage response, was of special interest to us among all the newly found interacting partners of the TTF1. We further performed co-immunoprecipitation (Co-IP) with DDB1 antibody and pulled down TTF1 confirming their interaction (Figure 2A and B).

**Figure 1:**
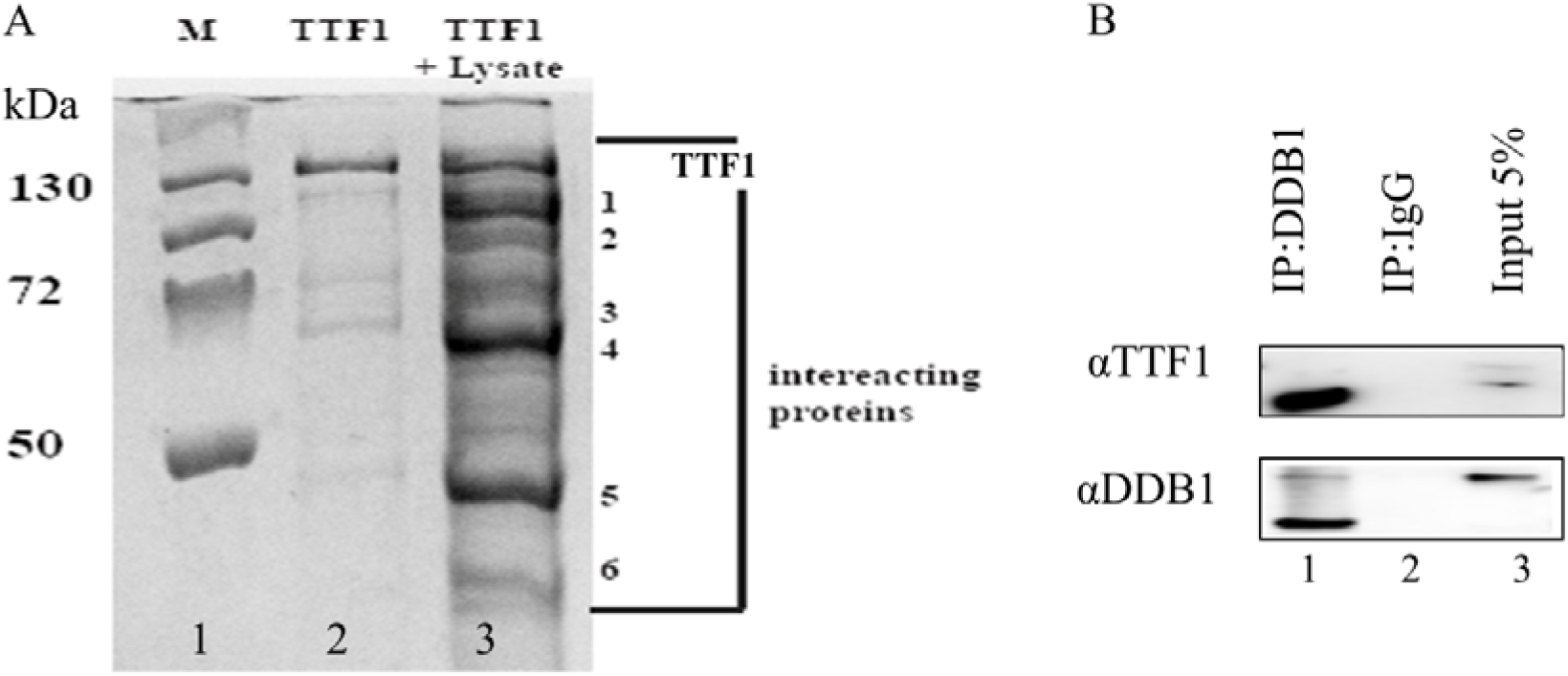
A) SDS PAGE gel showing Pull down using immobilized purified TTF1 protein. Lane 1 represents molecular weight marker. Lane 2 shows immobilized TTF1 only, Lane 3 represents eluate after HEK293T cell lysate incubated with bacterially purified TTF1 B) Co-Immunoprecipitation with DDB1 antibody; HEK293T cell lysate was either incubated with immobilized IgG (Lane 2) or DDB1 antibody (Lane 3), washed, eluted, and immunoblotted with respective antibodies as mentioned. IP represents 15% of the whole input.

### 2. TTF1 suppression downregulates DDB1

Since TTF1 is a transcription factor (TF), to investigate its role in regulation of DDB1 we silenced TTF1 gene (siTTF1) in HEK293T cells and observed the expression of DDB1 and viceversa. We discovered that TTF1 suppression also suppresses DDB1 protein level (Figure 3A and B) while DDB1 suppression elevated the TTF1 protein level (Figure 3C and D). Hence, expression of both the proteins are antagonistic to each other (Figure 3).

**Figure 3:**
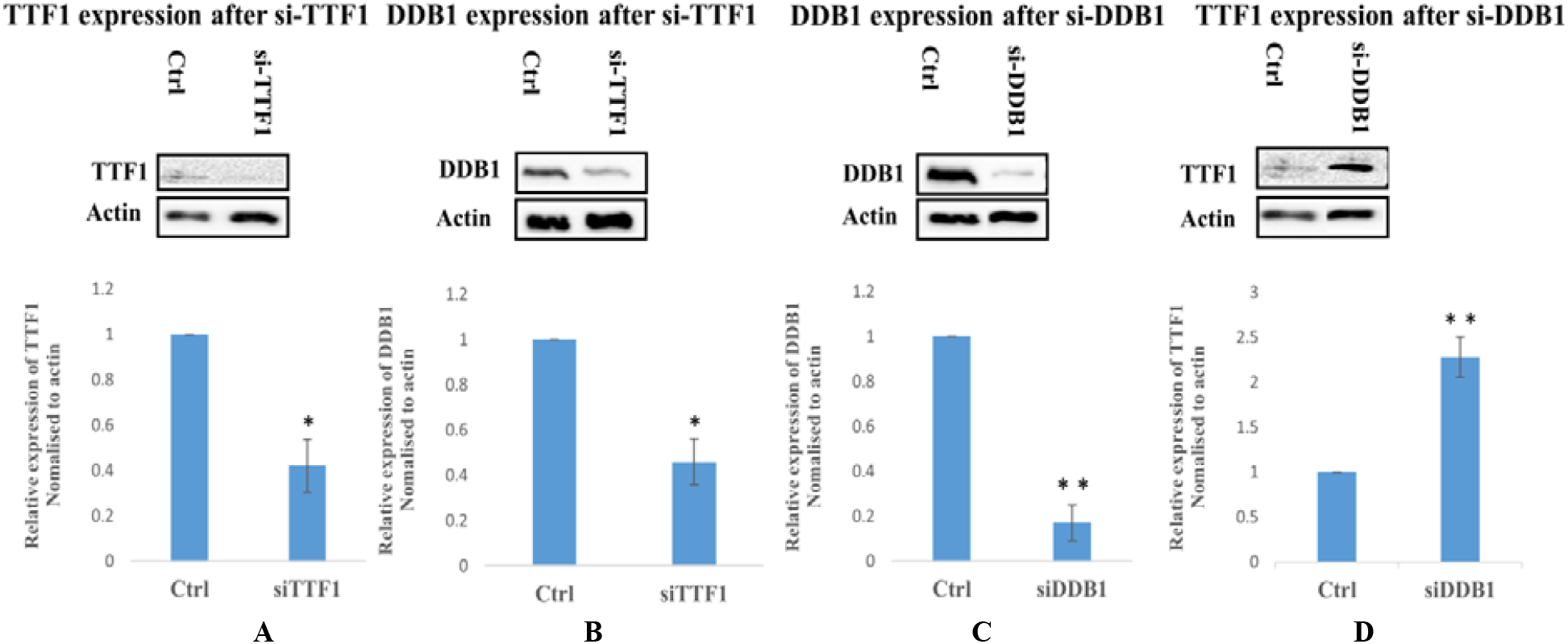
Western blot analysis after siRNA transfection in HEK293T cells; A, B) Expression profile of TTF1 and DDB1 respectively after Si-TTF1; C, D) Expression profile of DDB1 and TTF1 respectively after Si-DDB1 transfection. Bottom plots represent quantification by densitometric analysis (normalized to actin values). The mean values (± S.D.), obtained from 3 independent experiments are reported. Statistical analysis was performed by t test. *P < 0.05; **P < 0.005.

### 3. TTF1 is involved in UV mediated DNA damage repair response

Since TTF1 directly interacts with DDB1 and involved in its regulation, we further wanted to investigate the role of TTF1 in UV mediated DNA damage response. In order to explore the same, we studied the expression profile of TTF1 after 5 min, 10 min and 15 min post UV exposure. We observed that there is gradual increase in TTF1 protein concentration upto 15 mins post UV treatment (Figure 4). We also performed confocal microscopy to look for the localization of TTF1 at the UV-C induced DNA damaged sites using _γ_H2AX as a marker (Figure 5A Vs 5B). We found that concentration of TTF1 is considerably high at the DNA damaged site after 30 J/m^2^ of UV-C treatment to the HEK293T cells (Figure 5C Vs 5D) confirming the participation of TTF1 in UV mediated DNA damage response.

**Figure 4:**
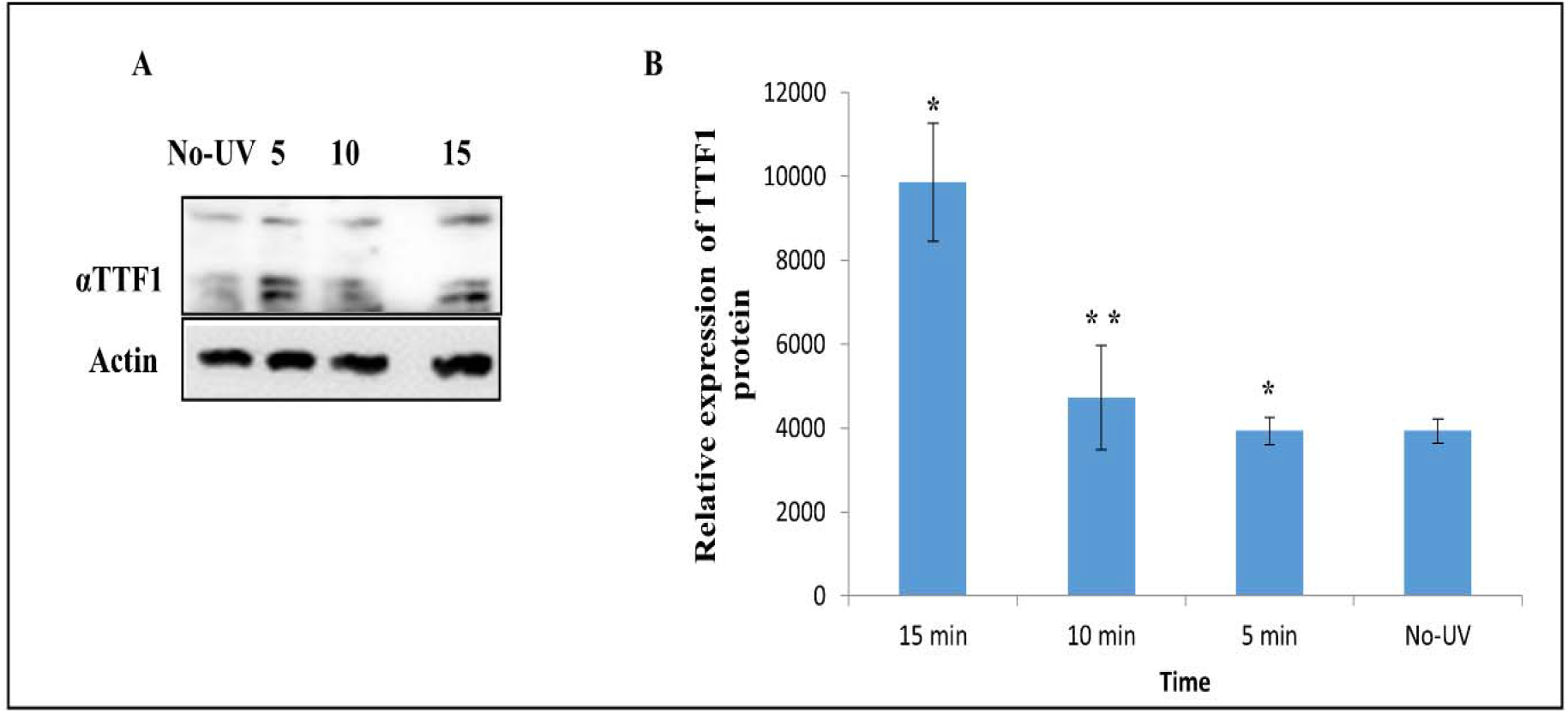
A) Western blot analysis of TTF1 after 30 J/m^2^ of UV irradiation at 5 min intervals; B) TTF1 protein levels in similar experiments, as those shown in (A), were quantified by densitometric analysis and normalized to actin values. The mean values (± S.D.), obtained from 3 independent experiments are reported. Statistical analysis was performed using t test. *P < 0.05; **P < 0.005.

**Figure 5:**
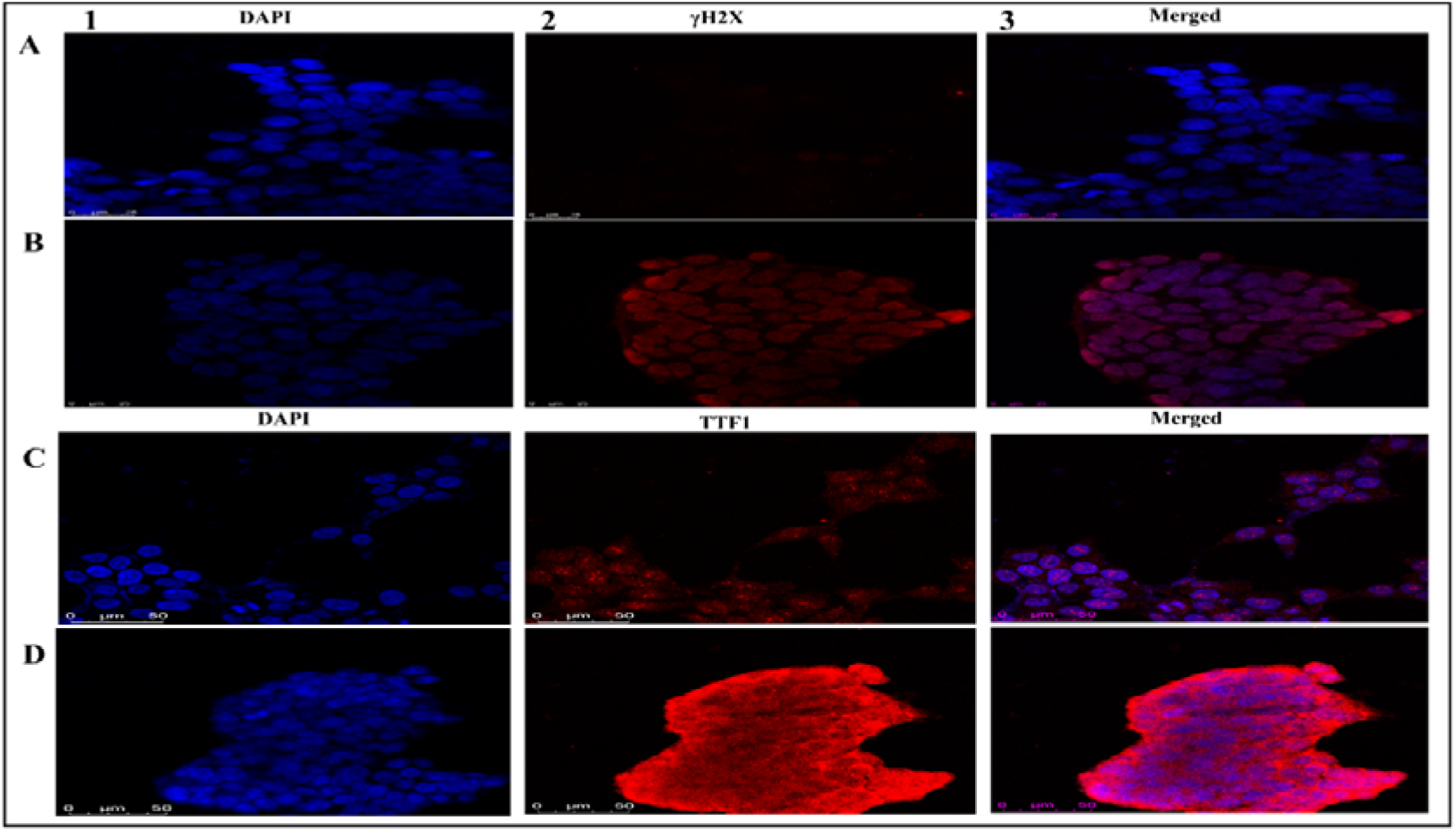
Confocal microscopy of HEK293T cells A) Expression of _γ_H2AX without (W/O) UV irradiation (panel 2A); B) γH2AX expression after 30 J/m^2^ UV irradiation (panel 2B); C) Expression of TTF1 W/O UV irradiation (panel 2C); D) TTF1 expression after 30 J/m^2^ UV irradiation (panel 2D). Panel 1 shows DAPI stained cells (for nuclear position) while panel 3 represents the merged images (panel 1 and panel 2).

## Discussion

Proper functioning of DNA repair mechanism is essential for maintenance of genome stability. Both sensing of DNA damage and repair (especially NER) are greatly influenced by TFs, particularly those TFs that bind to DNA at particular genomic loci. These factors are reported to regulate NER in both positive and negative ways [1,16,17]. Even after all these findings, the exact mechanism behind their activity is not very well understood. Mammalian TTF1 is an essential TF with multiple functions (as mentioned in introduction), which interacts with various proteins including CSB-B to cater the needs [11,13]. Our analysis of the result obtained through LC-MS confirmed various interactors reported earlier and also revealed several potential novel interacting partners such as DDB1 which are involved in DNA damage repair pathways and mutation in the same has been linked with various diseases including neurological and developmental process [17]. Our finding shows silencing of TTF1 downregulates DDB1 while silencing of DDB1 increases level of TTF1. It is possible that TTF1 goes through ubiquitination mediated proteasomal degradation by CRL4-DDB1 E3 ligase system. We further discovered that there is gradual increase and accumulation in the concentration of TTF1 after UV treatment of the HEK293T cells at the DNA damaged sites along with _γ_H2AX. These results suggest direct involvement of TTF1 in the UV induced DNA damge response and probably repair as well, which needs to be explored further for understanding the underlying mechanism. Also required is to decipher how exactly DDB1 regulates TTF1 activity at the damaged site. It’s already reported that DDB1 and TTF1 both interacts with CSA/B [5,11], hence it will be interesting to study what role these protein play to activate/regulate each other and what is the combined function etc. (Figure 6).

**Figure 6:**
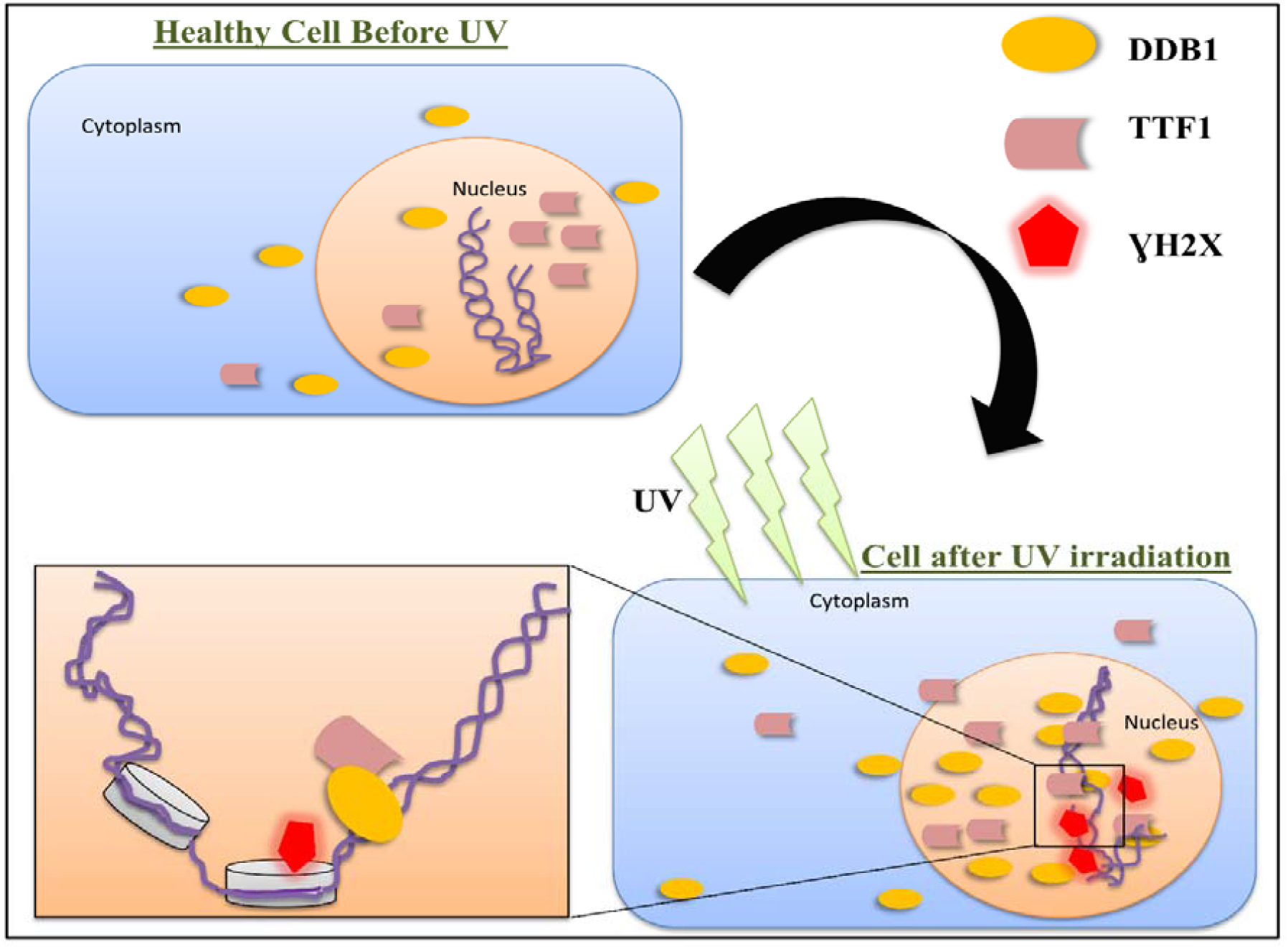
Proposed schematic representation of the role of TTF1 in UV-induced DNA damage sensing (and probably repair as well).

Our study is the first to report a novel function of this essential gene in DNA damage response, thereby opening up the possibility to use it as a therapeutic target in cancer drug formulation.

## Acknowledgement

Authors are thankful to the coordinator, School of Biotechnology, Dean and Director Institute of Science, Banaras Hindu University for providing space and facilities to conduct the research. We are highly thankful to Central Discovery Centre (CDC) and SATHI BHU for facilitating high resolution confocal microscope facility. We are also thankful to Advance Technology Platform Centre (ATPC), Regional Centre for Biotechnology, Faridabad for LCMS analysis. Further, we are thankful to Department of Biotechnology (DBT), Govt. of India for funding SKS with RLS grant (BT/RLF/Re-entry/43/2016), and JRF fellowship to KT. We are also thankful to IoE, BHU and University Grant Commission for providing funds to SKS.

## Supplementary information

**Figure S1:**
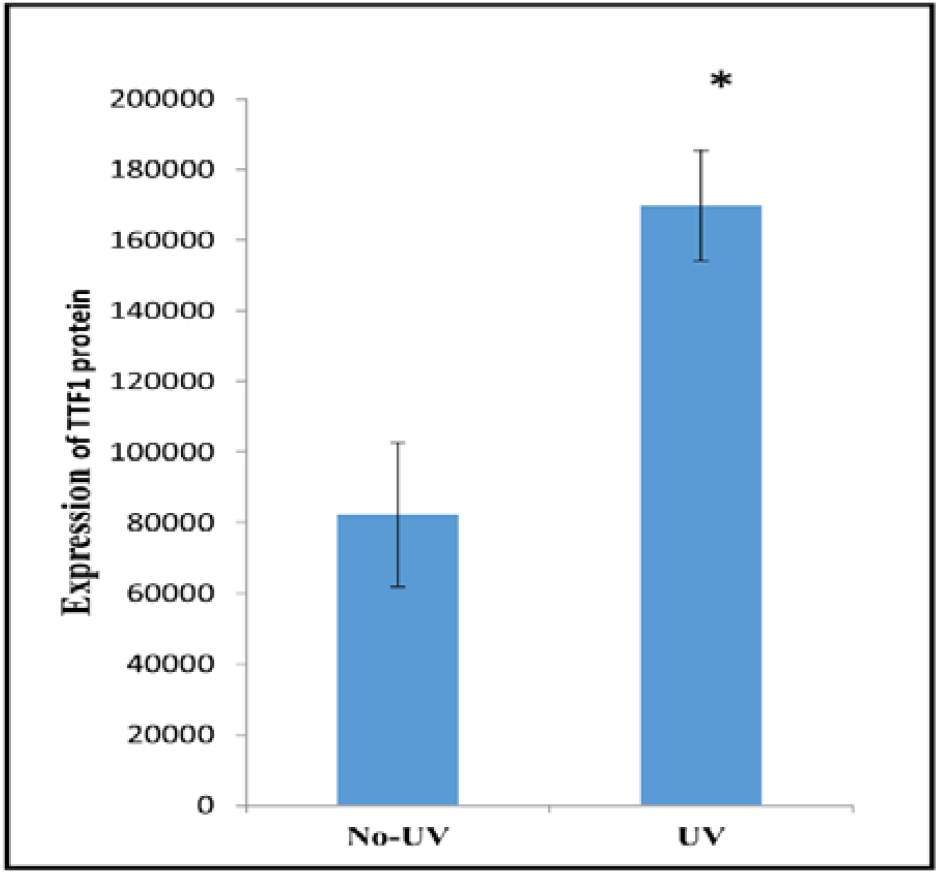
Quantification of Fluorescence intensity (based on Fig 6) by counting 5 different fields per sample. The mean values (± S.D.), obtained from 3 independent experiments are reported. Statistical analysis was performed using t test. *P < 0.05; **P < 0.005.

